# Functional selectivity for face processing in the temporal voice area of early deaf individuals

**DOI:** 10.1101/154138

**Authors:** Stefania Benetti, Markus J. van Ackeren, Giuseppe Rabini, Joshua Zonca, Valentina Foa, Francesca Baruffaldi, Mohamed Rezk, Francesco Pavani, Bruno Rossion, Olivier Collignon

## Abstract

Brain systems supporting face and voice processing both contribute to the extraction of important information for social interaction (e.g., person identity). How does the brain reorganize when one of these channels is absent? Here we explore this question by combining behavioral and multimodal neuroimaging measures (magneto-encephalography and functional imaging) in a group of early deaf humans. We show enhanced selective neural response for faces and for individual face coding in a specific region of the auditory cortex that is typically specialized for voice perception in hearing individuals. In this region, selectivity to face signals emerges early in the visual processing hierarchy, shortly following typical face-selective responses in the ventral visual pathway. Functional and effective connectivity analyses suggest reorganization in long-range connections from early visual areas to the face-selective temporal area in individuals with early and profound deafness. Altogether, these observations demonstrate that regions that typically specialize for voice processing in the hearing brain preferentially reorganize for face processing in born deaf people. Our results support the idea that cross-modal plasticity in case of early sensory deprivation relates to the original functional specialization of the reorganized brain regions.

## Introduction

The human brain is endowed by the fundamental ability to adapt its neural circuits in response to experience. Sensory deprivation has long been championed as a model to test how experience interacts with intrinsic constraints to shape functional brain organization. In particular, decades of neuroscientific research have gathered compelling evidence that blindness and deafness are associated with crossmodal recruitment of the sensory deprived cortices (1). For instance, in early deaf individuals, visual and tactile stimuli induce responses in regions of the cerebral cortex that are sensitive primarily to sounds in the typical hearing brain (2, 3).

Animal models of congenital and early deafness suggest that specific visual functions are relocated to discrete regions of the reorganized cortex and that this functional preference in cross-modal recruitment supports superior visual performance. For instance, superior visual motion detection is selectively altered in deaf cats when a portion of the dorsal auditory cortex, specialized for auditory motion processing in the hearing cat, is transiently deactivated (4). These results suggest that crossmodal plasticity associated with early auditory deprivation follows organizational principles that maintain the functional specialization of the colonized brain regions. In humans, however, there is only limited evidence that specific non-auditory inputs are differentially localized to discrete portions of the auditory-deprived cortices. For example, Bola and colleagues have recently reported, in deaf individuals, crossmodal activations for visual rhythm discrimination in the posterior-lateral and associative auditory regions that are recruited by auditory rhythm discrimination in hearing individuals (5). However, the observed cross-modal recruitment encompassed an extended portion of these temporal regions, which were found activated also by other visual and somatosensory stimuli and tasks in previous studies (2,3). Moreover, it remains unclear whether specific reorganization of the auditory cortex contributes to the superior visual abilities documented in the early deaf humans (6). These issues are of translational relevance since auditory re-afferentation in the deaf is now possible through cochlear implants and cross-modal recruitment of the temporal cortex is argued to be partly responsible for the high variability in speech comprehension and literacy outcomes (7), which still poses major clinical challenges.

To address these issues, we tested whether, in early deaf individuals, face perception selectively recruits discrete regions of the temporal cortex that typically respond to voices in hearing people. Moreover, we explored if such putative face-selective cross-modal recruitment is related to superior face perception in the early deaf. We used face perception as a model based on its high relevant social and linguistic valence for deaf individuals and the suggestion that auditory deprivation might be associated with superior face processing abilities (8). Recently, it was demonstrated that both linguistic (9) and non-linguistic (10) facial information remap to temporal regions in postlingually deaf individuals. In early deaf individuals, we expected to find face-selective responses in the middle and ventro-lateral portion of the auditory cortex, a region showing high sensitivity to vocal acoustic information in hearing individuals, namely the “temporal voice-selective area” (TVA)(11). This hypothesis is notably based on the observation that facial and vocal signals are integrated in lateral belt regions of the monkey temporal cortex (12). Moreover, there is evidence for functional interactions between this portion of the TVA and the face-selective area of the ventral visual stream in the middle lateral fusiform gyrus (the fusiform face area, FFA)(13) during person recognition in hearing individuals (14), and of direct structural connections between these regions in hearing individuals (15). In order to further characterize the potential role of reorganized temporal cortical regions in face perception, we also investigated whether these regions support face identity discrimination by means of a repetition-suppression experiment in functional magnetic resonance imaging (16). Next, we investigated the time-course of putative TVA activation during face perception by reconstructing virtual time-series from MEG recordings while subjects viewed images of faces and houses. We predicted that, if deaf TVA has an active role in face perception, category-selectivity should be observed close in time to the first selective response to faces in the fusiform gyrus, i.e. between 100-200ms (17). Finally, we examined the role of long-range cortico-cortical functional connectivity in mediating the potential cross-modal reorganization of TVA in the deaf.

## Results

### Experiment 1: Face perception selectively recruits rTVA in early deaf compared to hearing individuals

To test whether face perception specifically recruits auditory voice-selective temporal regions in the deaf group (= 15) we functionally localized (i) the TVA in a group of hearing controls (=15) with an fMRI voice localizer and (ii) the face-selective network in each group (i.e. hearing controls = 16; hearing users of the Italian Sign Language = 15; and deaf individuals = 15) with a fMRI face localizer contrasting full-front images of faces and houses matched for low-level features like color, contrast and spatial frequencies (see experimental procedures). A group of hearing users of the Italian Sign Language (LIS) was included in the experiment to control for the potential confounding effect of exposure to visual language. Consistent with previous studies of face (13) and voice (11) perception, face-selective responses were observed primarily in the mid-lateral fusiform gyri bilaterally as well as in the right posterior superior temporal sulcus (pSTS) across the three groups (Fig. S1 and Table 1) while voice-selective responses were observed in the mid-lateral portion of the superior temporal gyrus (mid-STG) and the mid-upper bank of the STS (mid-STS) in the hearing control group (Fig. S2).

**Table 1.** Regional responses for the main effect of face condition in each group and differences between the three groups.

| Area | Cluster size | X<sub>(mm)</sub> | Y<sub>(mm)</sub> | Z<sub>(mm)</sub> | Z | D.F. | P<sub>FWE</sub> |
| --- | --- | --- | --- | --- | --- | --- | --- |
| *Hearing Controls Faces > Houses* |  |  |  |  |  | 15 |  |
| R fusiform gyrus (lateral) | 374 | 44 | -50 | -16 | 4.89 |  | 0.017* |
| R sup. temporal gyrus/sulcus (post.) | 809 | 52 | -42 | 16 | 4.74 |  | < 0.001 |
| R middle frontal gyrus | 1758 | 44 | 8 | 30 | 4.33 |  | < 0.001 |
| L fusiform gyrus (lateral) | 97 | -42 | -52 | -20 | 4.29 |  | 0.498 |
| *Hearing-LIS Faces > Houses* |  |  |  |  |  | 14 |  |
| R sup. temporal gyrus/sulcus (post.) | 1665 | 52 | -44 | 10 | 5.64 |  | < 0.001* |
| R middle frontal gyrus | 3596 | 34 | 4 | 44 | 5.48 |  | < 0.001* |
| R inferior frontal gyrus | s.c. | 48 | 14 | 32 | 5.45 |  |  |
| R inferior parietal gyrus | 1659 | 36 | -52 | 48 | 5.45 |  | < 0.001* |
| R fusiform gyrus (lateral) | 46^ | 44 | -52 | -18 | 3.52 |  | 0.826 |
| L middle frontal gyrus | 1013 | -40 | 4 | 40 | 5.13 |  | < 0.001* |
| L fusiform gyrus (lateral) | 120 | -40 | -46 | -18 | 4.02 |  | 0.282 |
| R/L superior frontal gyrus | 728 | 2 | 20 | 52 | 4.78 |  | < 0.001 |
| R/L superior frontal gyrus | 728 | 2 | 20 | 52 | 4.78 |  | < 0.001 |
| *Deaf Faces > Houses* |  |  |  |  |  | 14 |  |
| R middle frontal gyrus | 1772 | 42 | 2 | 28 | 4.84 |  | < 0.001* |
| R inferior temporal gyrus | 845 | 50 | -60 | -10 | 4.04 |  | < 0.001 |
| R fusiform gyrus (lateral) | s.c. | 48 | -56 | -18 | 3.54 |  |  |
| R sup. temporal gyrus/sulcus (post.) | s.c. | 50 | -40 | 14 | 4.01 |  |  |
| R sup. temporal gyrus/sulcus (mid.) | 64 | 54 | -24 | -4 | 3.80 |  | 0.005° |
| R thalamus (posterior) | 245 | 10 | -24 | 10 | 4.02 |  | 0.042 |
| L fusiform gyrus (lateral) | 209 | -40 | -66 | -18 | 3.90 |  | 0.071 |
| R putamen | 329 | 28 | 0 | 4 | 4.49 |  | 0.013 |
| L middle frontal gyrus | 420 | -44 | 26 | 30 | 3.95 |  | 0.004 |
| *Deaf > Hearing Controls ∩ Hearing-LIS Faces > Houses* |  |  |  |  |  | 3,44 |  |
| R sup. temporal gyrus/sulcus (middle) | 73 | 62 | -18 | 2 | 3.86 |  | 0.006° |
| *Deaf > Hearing Controls Faces > Houses* |  |  |  |  |  | 30 |  |
| R sup. temporal gyrus/sulcus (middle) | 167 | 62 | -18 | 4 | 3.77 |  | 0.001° |
| L sup. temporal gyrus/sulcus (middle) | 60 | -64 | -24 | 10 | 3.64 |  | 0.007° |
| *Deaf > Hearing-LIS - Faces > Houses* |  |  |  |  |  | 29 |  |
| R sup. temporal gyrus/sulcus (middle) | 73 | 62 | -18 | 2 | 3.86 |  | 0.006° |

Significance corrections are reported at the cluster level; cluster size threshold = 50;(^) cluster size < 50; (\*) brain activations significant after FWE voxel-correction over the whole brain; (°) brain activation significant after FWE voxel-correction over a small spherical volume (25mm radius) at peak-coordinates for right and left hearing TVA. Abbreviations: FWE, Family-Wise Error, L, Left; R, Right; D.F., Degrees of Freedom; s.c., same cluster.

When selective neural responses to face perception were compared between the three groups, enhanced face selectivity was observed in the right mid-lateral STG extending ventrally to the mid-upper bank of the STS (MNI coordinates [62 −18 2]) in the deaf group compared to both the hearing and the hearing-LIS groups (Fig. 1A-B; Table 1). The location of this selective response strikingly overlapped with the superior portion of the right TVA as functionally defined in our hearing control group (Fig. 1C-D). Face selectivity was additionally observed in the left dorsal STG posterior to TVA (MNI coordinates [−64 −28 8]) when the deaf and hearing control group where compared; however, no differences were detected in this regions when the deaf and hearing control groups were, respectively, compared to hearing-LIS users. In order to further describe the preferential face response observed in the right temporal cortex, we extracted individual measures of estimated activity (beta weights) in response to faces and houses from the right TVA as independently localized in the hearing groups. In these regions, an analysis of variance revealed an interaction effect (F_[Category × Group]=_ 16.18, p < 0.001, η^2^ = 0.269) confirming increased face-selective response in the right mid-STG/STS of deaf individuals compared to both the hearing controls and hearing LIS users (t_[deaf > hearing]_ = 3.996, p < 0.001;, Cohen’s-d = 1.436; t_[deaf>hearing-LIS]_ = 3.907, p < 0.001, Cohen’s-d = 7.549; Fig. 1D). Although no face selectivity was revealed - at the whole brain level and with small volume correction (SVC) - in the left temporal cortex of deaf individuals, we further explored the individual responses in left mid-TVA for completeness. Cross-modal face selectivity was also revealed in this region in the deaf, albeit the inter-individual variability within this group was larger and the face-selective response was weaker (see supplemental information and Fig. S4). In contrast to the preferential response observed for faces, no temporal region showed group-differences for house-selective responses (Table 1; Fig. S1). Hereafter, we focus on the right temporal region showing robust face-selective recruitment in the deaf and refer to it as the deaf Temporal Face Area (dTFA).

**Figure 1.**
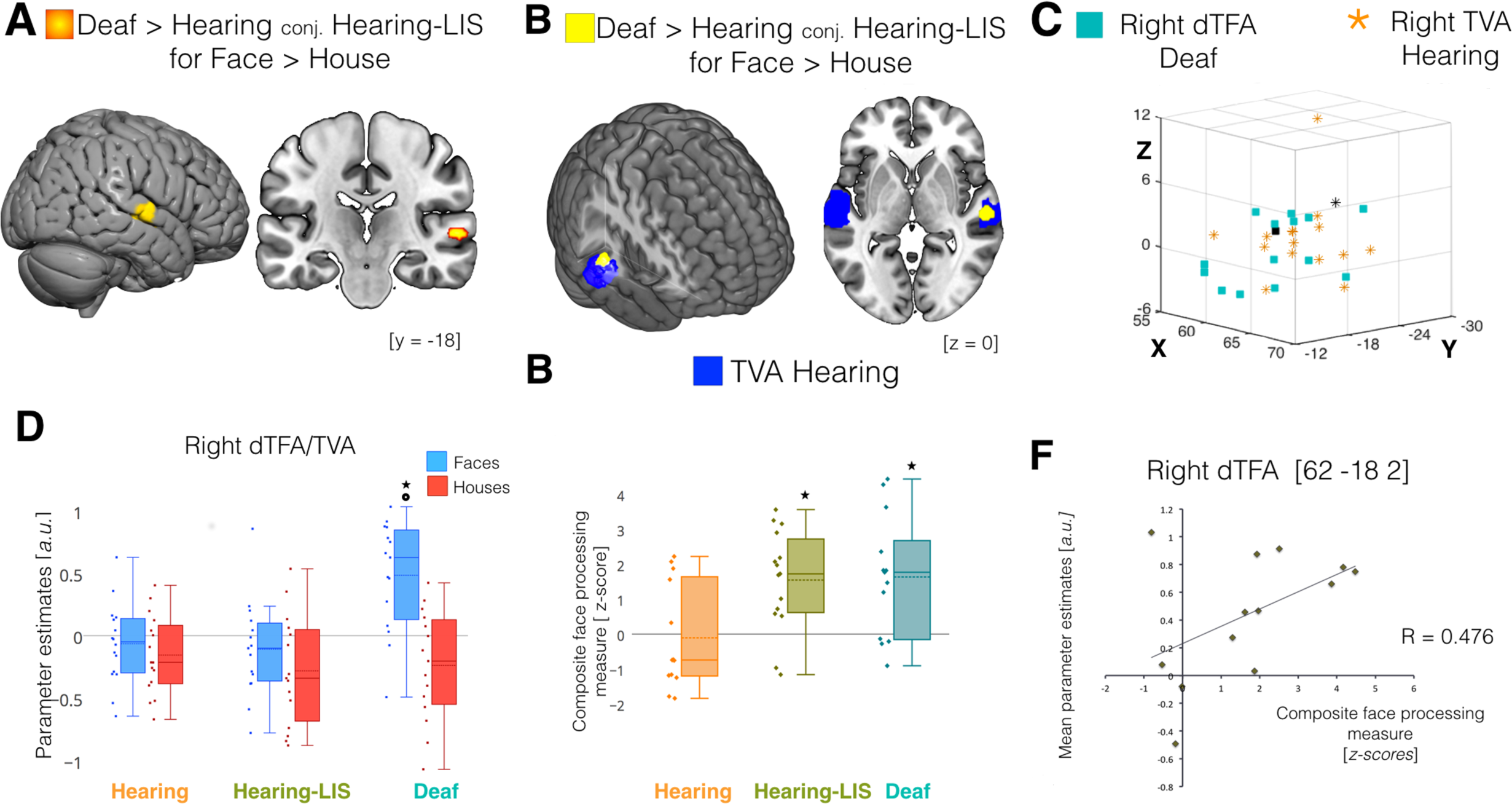
Cross-modal recruitment of right dTFA in the deaf. Regional responses significantly differing between groups during face compared to house processing are depicted over multi-planar slices and renders of the MNI-ICBM152 template. **(A)** Supra-threshold cluster (P<0.05 FWE small volume-corrected) showing difference between deaf subjects compared to both hearing subjects and hearing LIS users (conj.= conjunction analysis). **(B)** Depiction of the spatial overlap between face-selective response in deaf subjects (yellow) and the voice-selective response in hearing subjects (blue) in the right hemisphere. **(C)** 3D scatterplot depicting individual activation peaks in mid STG/STS for faces selective responses in early deaf subjects (cyan squares) and voices selective responses in hearing subjects (orange stars); black markers represent the group-maxima for face-selectivity in the right DTFA of deaf subjects (square) and voice-selectivity in right TVA of hearing subjects (star). **(D)** Box-plots showing the central tendency (*a. u.,* arbitrary unit; median = solid line; mean = dashed line) of activity estimates for face (blue) and house (red) processing computed over individual parameters (diamonds) extracted at group-maxima for rTVA in each group; * P<0.001 between groups; **°** P<0.001 for Faces > Houses in deaf subjects. **(E)** Box-plots showing central tendency for composite face processing scores (*z-scores*; solid line = median; dashed line = mean) for the three groups; * P < 0.016 for deaf > hearing and hearing-LIS > hearing. (**F)** Scatterplot displaying a trend for significant positive correlation (P=0.05) between individual face-selective activity estimates and composite measures of face processing ability in deaf subjects.

At the behavioral level, performance in a well-known and validated neuropsychological tests of individual face matching, the Benton Facial Recognition Test (18) and a delayed recognition of facial identities seen in the scanner were combined in a composite face-recognition measure in each group. This composite score was computed in order to achieve a more stable and comprehensive measure of the underlying face processing abilities (19).When the three groups were compared on face processing ability, the deaf group significantly outperformed the hearing group (t = 3.066, p = 0.012, Cohen’s-d = 1.048; Fig. 1E) but not the hearing-LIS group, which also performed better than the hearing group (t = 3.080, p = 0.011, Cohen’s-d = 1.179; Fig. 1E). This is consistent with previous observations suggesting that both auditory deprivation and use of sign language lead to superior ability to process face information (20). To determine whether there was a relationship between face-selective recruitment of the dTFA and face perception we compared inter-individual differences in face-selective responses with corresponding variations on the composite measure of face recognition in deaf individuals. Face-selective responses in the right dTFA showed a trend for significant positive correlation with face processing performance in the deaf group (R_deaf_ = 0.476, CI = [- 0.101 0.813], p = 0.050; Fig. 1E). Neither control group showed a similar relationship in the right TVA (R_hearing subjects_ = 0.038, CI = [-0.527 0.57], p = 0.451; R_hearing-LIS_ = −0.053, CI = [-0.55 0.472], p = 0.851). No significant correlation was detected between neural and behavioral responses to house information deaf subject (R=0.043, p = 0.884). Moreover, behavioral performances for the house and face tests did not correlate with LIS exposure. It is however important to note that the absence of a significant difference in strength of correlation between deaf and hearing groups (see confidence intervals reported above) limits our support for the position that crossmodal reorganization is specifically linked to face perception performance in deaf individuals.

### Experiment 2: Reorganized right dTFA codes individual face identities

To further evaluate whether reorganized dTFA is also able to differentiate between individual faces we implemented a second experiment using fMR-adaptation (16). Recent studies in hearing individuals have found that a rapid presentation rate, with a peak at about 6 face stimuli by second (6 Hz), leads to the largest fMRI-adaptation effect in ventral occipito-temporal face-selective regions, including the FFA, indicating optimal individualization of faces at these frequency rates (21, 22). Participants were presented with blocks of identical or different faces at five frequency rates of presentation between 4 and 8.5 Hz. Individual beta values were estimated for each condition (same/different faces × 5 frequencies) individually in the right FFA (in all groups), TVA (in hearing subjects and hearing LIS users) and dTFA (in deaf subjects).

Since there were no significant interactions in the TVA ([group] × [identity] × [frequency], p = 0.585) or FFA ([group] × [identity] × [frequency], p = 0.736) or group effects (TVA, p=0.792; FFA, p=0.656) when comparing the hearing and hearing-LIS groups, they were merged in a single group for subsequent analyses. With the exception of a main effect of *Face Identity*, reflecting the larger response to different than identical faces for deaf and hearing participants (Fig. 2B), there were no other significant main or interaction effects in the right FFA. In the TVA/dTFA clusters, in addition to a main effect of *Face identity* (p < 0.001), we also observed two significant interactions of [group] × [face identity] (p = 0.013) and of [group] × [identity] × [frequency] (p = 0.008). A post-hoc *t*-test revealed a larger response to different faces (p = 0.034) across all frequencies in deaf compared to hearing participants. In addition, the significant three-way interaction was driven by larger responses to different faces between 4 and 6.6 Hz (4 Hz: p = 0.039; 6 Hz: p= 0.039; 6.6 Hz: p = 0.003; Fig. 2A) in deaf compared to hearing participants. In this averaged frequency range, there was a trend for significant release from adaptation in hearing participants (p =0.031; for this test the significance threshold was *p* = 0.05/2 groups = 0.025) and a highly significant effect of release in deaf subjects (p < 0.001); when the two groups were directly compared, the deaf group also showed larger release from adaptation compared to hearing and hearing-LIS participants (p <0.001; Fig. 2B). These observations reveal not only that the right dTFA shows enhanced coding of individual face identity in deaf individuals but also suggest that the right TVA may show a similar potential in hearing individuals.

**Figure 2.**
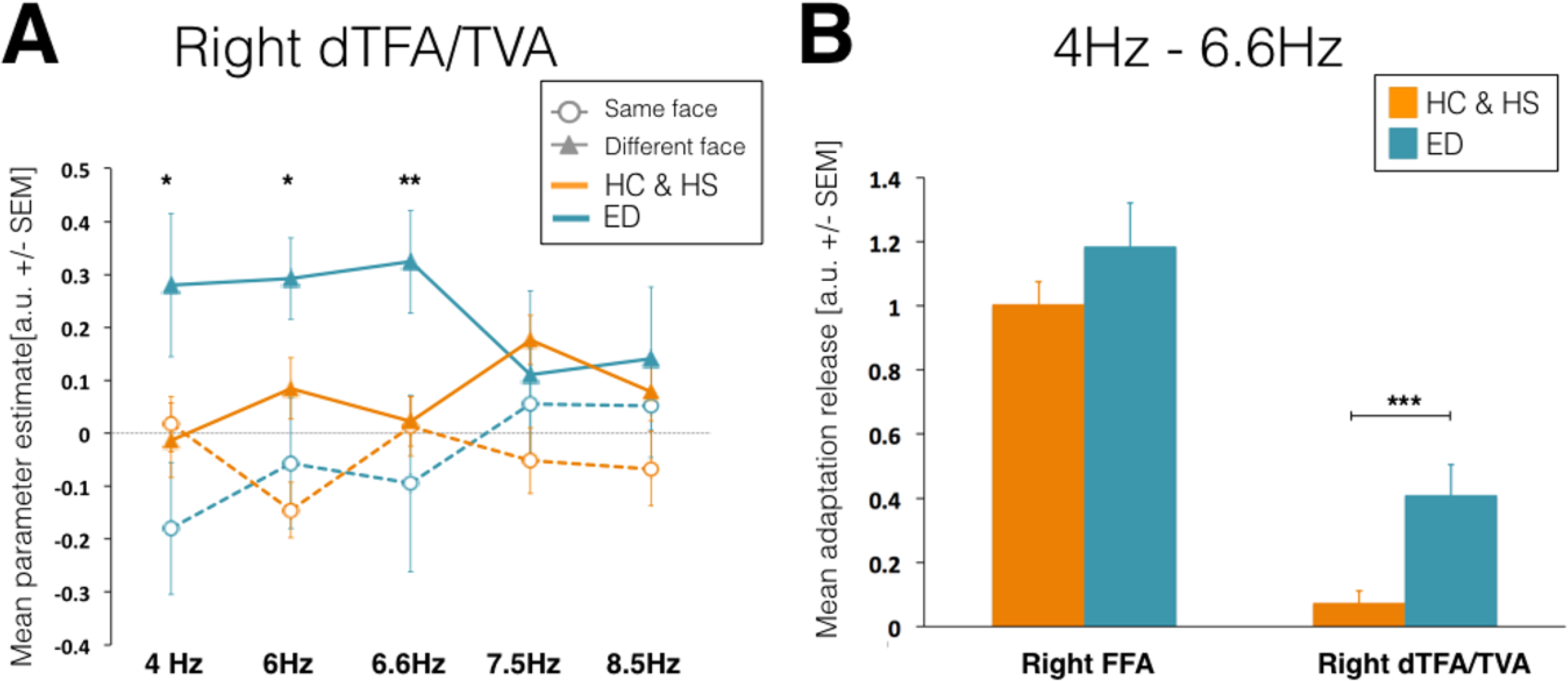
Adaptation to face identity repetition in the right dTFA of deaf individuals. . **(A)** Mean activity estimates (beta weights; *a. u.* ± SEM) are reported at each frequency rate of stimulation for same (empty circle – dashed line) and different (full triangle – solid line) faces in both deaf (cyan) and hearing (orange) individuals. Deaf participants show larger responses for different faces at 4 to 6.6 Hz (\**P* <0.05; \*\**P* < 0.01) compared to hearing individuals. **(B)** Bar graphs show the mean adaptation-release estimates (*a.u.* ± SEM) across frequencies rates 4 to 6.6 Hz in the right FFA and right dTFA/TVA in deaf (orange) and hearing (cyan) individuals. In deaf subjects the release from adaptation to different faces is above baseline (*P <* 0.001) and larger than in hearing individuals (\*\*\**P* < 0.001). No significant differences are found in right FFA. ED, early deaf; HC, hearing controls; HS, hearing-LIS controls.

### Experiment 3: Early selectivity for faces in right dTFA

In a third neuroimaging experiment, magneto-encephalographic (MEG) responses were recorded during an oddball task with the same face and house images used in the fMRI face localizer. Since no differences were observed between the hearing and hearing-LIS groups for the fMRI face localizer experiment, only deaf subjects (=17) and hearing (=14) participants were included in this MEG experiment.

Sensor-space analysis on evoked responses to Face and House stimuli was performed using permutation statistics, and corrected for multiple comparisons with a maximum cluster-mass threshold. Clustering was performed across space (sensors), and time (100-300ms). Robust face selective responses across groups (*p*<.005, cluster-corrected) were revealed in a large number of sensors mostly around 160-210 ms (Fig 3A) in line with previous observations (23). Subsequent time domain beamforming (LCMV) on this time window of interest showed face selective regions of the classical face-selective network, including the FFA (Fig 3B-C, top panel). To test whether dTFA, as identified in fMRI, is already recruited during this early time-window of face perception we tested whether face selectivity was higher in the deaf versus hearing group. For increased statistical sensitivity, a small volume correction was applied using a 15mm sphere around the voice-selective peak of activation observed in the hearing group in fMRI (MNI x = 63; y = −22; z = 4). Independently reproducing our fMRI results, we observed enhanced selective responses to faces versus houses in deaf when compared to hearing subjects specifically in the right middle temporal gyrus (Fig. 3B-C, bottom panel).

**Figure 3.**
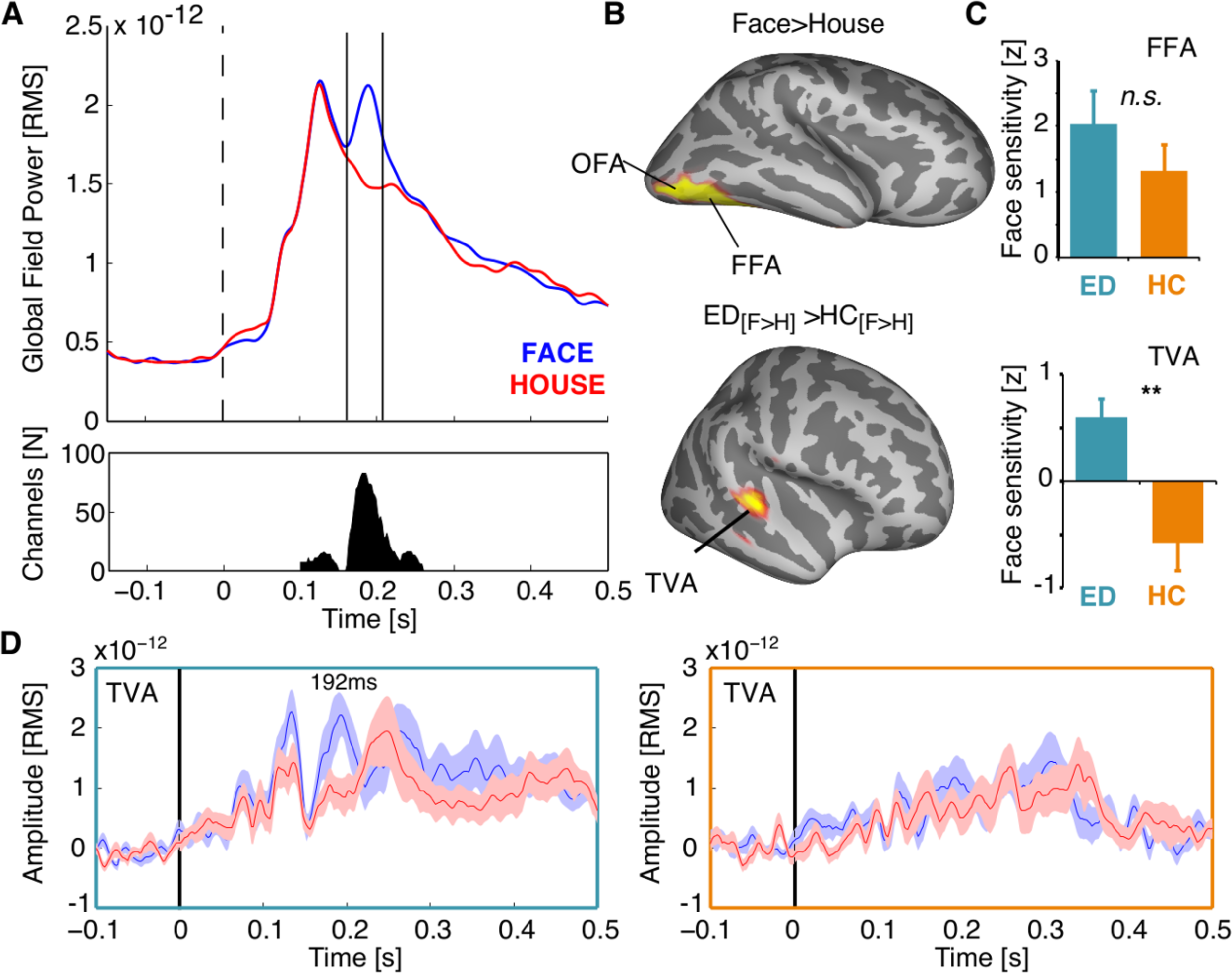
Face selectivity in right dTFA is observed within 200ms post-stimulus. (**A)** Global field power of the evoked response for faces (blue) and houses (red) across participants. The bottom panel depicts the number of sensors contributing to the difference between the two conditions (P<.005, cluster-corrected) at different points in time. Vertical bars in the top panel mark the time window of interest (160-210ms) for source reconstruction. **(B)** The top panel shows face selective regions within the time window of interest in ventral visual areas across groups (P<.05, FWE). The bottom panel highlights the interaction effect between groups (P<.05, FWE). **(C)** Bar graphs illustrate broadband face sensitivity (faces versus houses) for deaf (cyan) and hearing subjects (orange) at peak locations in FFA and TVA. An interaction effect is observed in dTFA (P<.005), but not FFA. N.S., not significant; (**), p<0.005. **(D)** Virtual sensors from the TVA peak locations show the averaged RMS time course for Faces and Houses in the deaf (left, cyan) and hearing (right, orange) group. Shading reflects the standard error of the mean (SEM). Face selectivity in the deaf group peaks at 192 ms in TVA of deaf individuals. No discernible peak is visible in TVA of the hearing group. Abbreviations: HC, Hearing Controls; ED = Early Deaf; OFA, Occipital Face Area.

Finally, to explore the timing of face selectivity in dTFA, virtual sensor time-courses were extracted for each group and condition from grid points close to the fMRI peak locations showing face- (FFA: hearing&deaf) and voice-selectivity (TVA: hearing subjects). We found a face-selective component in dTFA with a peak at 192ms (Fig. 3D), 16ms after the FFA peak at 176ms (Fig. 3D). In contrast, no difference between conditions is seen at the analogous location in the hearing group (Fig 3D).

### Long-range connections from V2/V3 support face-selective response in deaf TVA

Previous human and animal studies have suggested that long-range connections with preserved sensory cortices might sustain cross-modal reorganization of sensory deprived cortices (24). We first addressed this question by identifying candidate areas for the source of cross-modal information in right dTVA; to this end, a Psychophysiological Interactions (PPI) analysis was implemented and the face-selective functional connectivity between right TVA/dTFA and any other brain regions was explored. During face processing specifically, right dTFA showed a significant increase of inter-regional coupling with occipital and fusiform regions in the face-selective network extending to earlier visual associative areas in the lateral occipital cortex (V2/V3) of deaf individuals only (Fig. 4A). Indeed, when face-selective functional connectivity was compared across groups the effect that differentiated most strongly between deaf and both hearing and hearing-LIS individuals was in the right mid-lateral occipital gyrus (peak coordinates: x = 42, y = −86, z = 8; z = 5.91, cluster size = 1561, p < 0.001 FWE cluster- and voxel-corrected, fig. 4A and table S5). To further characterize the causal mechanisms and dynamics of the pattern of connectivity observed in the deaf group, we investigated effective connectivity to right dTFA by combining Dynamic Causal Modeling (DCM) and Bayesian Model Selection (BMS) in this group. Three different, neurobiologically plausible models were defined based on our observations and previous studies of face-selective effective connectivity in hearing individuals (25): the first model assumed that face-selective response in right dTFA was supported by increased direct ‘feed-forward’ connectivity from early visual occipital regions (right V2/V3); the two alternative models assumed that increased ‘feed-back’ connectivity from ventral visual face regions (right FFA) or posterior temporal face regions (right pSTS), respectively, would drive face-selective responses in right dTFA (Fig. 4B). While the latter two models showed no significant contributions, the first model, including direct connections from right V2/V3 to right TFA, accounted well for face-selective responses in this region of deaf individuals (exceedance probability = 0.815) (Fig. 4C).

**Figure 4.**
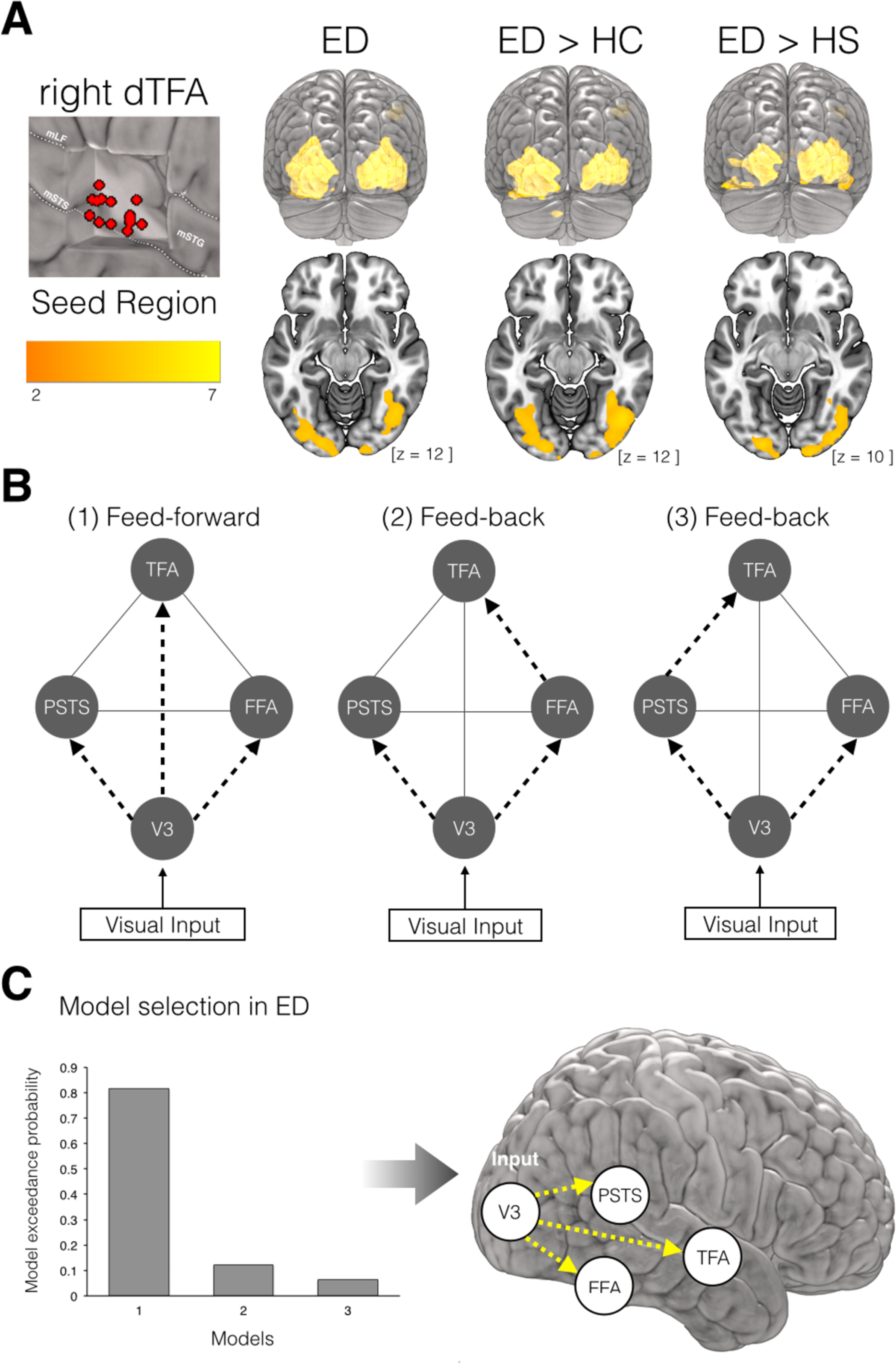
Functional and effective connectivity during face processing in early deaf. (**A**) Psycho-physiological interactions (PPI) seeding the right TVA/dTFA. In the left panel, individual loci of time-series extraction are depicted in red over a cut-out of the right mid STG/STS showing variability in peak of activation within this region in the deaf group; m = middle, LF = Lateral Fissure. In the right panel, supra-threshold (P = 0.05 FWE cluster-corrected over the whole brain) face-dependent PPI of rTVA in deaf subjects and significant differences between the deaf and the two control groups are superimposed on the MNI-ICBM152 template. (**B**) The three dynamic causal models (DCMs) used for the study of face-specific effective connectivity in the right hemisphere. Each model equally comprises: experimental visual inputs in V3, exogenous connections between regions (grey solid lines), and face-specific modulatory connections to FFA and pSTS (black dashed arrows). The three models differ in terms of the face-specific modulatory connections to dTFA. (**C**) Bayesian model selection showed that a modulatory effect of face from V3 to dTFA best fit the face-selective response observed in the deaf dTFA (left panel) as depicted in the schematic representation of face-specific information flow (right panel).

## Discussion

In this study we combined state-of-the-art multimodal neuroimaging and psychophysical protocols to unravel how early auditory deprivation triggers specific reorganization of auditory-deprived cortical areas to support the visual processing of faces. In deaf individuals, we report enhanced selective responses to faces in a portion of the mid-STS in the right hemisphere, a region overlapping with the right mid-TVA in hearing individuals (26) and that we refer to as the ‘deaf Temporal Face Area’. The magnitude of right dTFA recruitment in the deaf subjects showed a trend towards positive correlation with measures of individual face recognition ability in this group. Furthermore, significant increase of neural activity for different faces compared to identical faces supports individual face discrimination in the right dTFA of the deaf subjects. Using MEG, we found that face-selectivity in right dTFA emerges within the first 200ms following face onset, only slightly later than right FFA activation. Finally, we found that increased long-range connectivity from early visual areas best explained the face-selective response observed in the dTFA of deaf individuals.

Our findings add novelty to the observation of task-specific cross-modal recruitment of associative auditory regions reported by Bola and colleagues (5): to our knowledge, it is the first observation, in early deaf humans, of selective cross-modal recruitment of a discrete portion of the auditory cortex for specific and high-level visual processes typically supported by the ventral visual stream in the hearing brain. Additionally, we provide evidence for a functional relationship between recruitment of discrete portions of the auditory cortex and specific perceptual improvements in deaf individuals. The face-selective cross-modal recruitment of dTFA suggests that cross-modal effects does not occur uniformly across areas of the deaf cortex and supports the notion that cross-modal plasticity is related to the original functional specialization of the colonized brain regions(4, 27). Indeed, temporal voice areas typically involved in an acoustic-based representation of voice identity (28) are shown here to code for facial identity discrimination (see Fig 2A). This is in line with, previous investigations in blind humans, which have reported that cross-modal recruitment of specific occipital regions by non-visual inputs follows organizational principles similar to those observed in the sighted. For instance, following early blindness, the lexico-graphic components of Braille reading elicit specific activations in a left ventral fusiform region that typically responds to visual words in sighted individuals (29) while auditory motion selectively activate regions typically selective for visual motion in the sighted (30).

Crossmodal recruitment of a sensory-deprived region might find a “neuronal niche” in a set of circuits that perform functions that are sufficiently close to the ones required by the remaining senses (31). It is, therefore, expected that not all visual functions will be equally amenable to reorganization following auditory deprivation. Accordingly, functions targeting (supramodal) processes that can be shared across sensory systems (32, 33) or benefit from multisensory integration will be the most susceptible to selectively recruit specialized temporal regions deprived of their auditory input (4, 27). Our findings support this hypothesis since the processing of faces and voices share several common functional features, like inferring the identity, the affective states, the sex, the age of someone. Along those lines, no selective activity to houses was observed in the temporal cortex of deaf subjects, potentially due to the absence of a common computational ground between audition and vision for this class of stimuli. In hearing individuals, face-voice integration is central to person identity decoding (34), occurs in voice selective regions (35), and might rely on direct anatomical connections between the voice and face networks in the right hemisphere (15). Our observation of stronger face-selective activations in the right than left mid-STG/STS in deaf individuals further reinforces the notion of functional selectivity in the sensory-deprived cortices. In fact, similarly to face perception in the visual domain, the right mid-anterior STS regions respond more strongly than the left side to non-linguistic aspects of voice perception and contributes to the perception of individual identity, gender, age and emotional state by decoding invariant and dynamic voice features in hearing subjects (34). Moreover, our observation that right dTFA, similarly to right FFA, shows fMRI adaptation in response to identical faces, suggests that this region is able to process face identity information. This observation is also comparable with previous findings showing fMRI adaptation to speaker voice identity in right TVA of hearing individuals (36). In contrast, the observation of face selectivity in the posterior STG for deaf compared to hearing controls, but not hearing-LIS users, support the hypothesis that regions devoted to speech and multimodal processing in the posterior left temporal cortex might, at least in part, reorganize to process visual aspects of sign language (37).

We know from neurodevelopmental studies that, following an initial period of exuberant synaptic proliferation, projections between the auditory and visual cortices are eliminated either through cell death or retraction of exuberant collaterals during the synaptic pruning phase. The elimination of weaker, unused or redundant synapses is thought to mediate the specification of functional and modular neuronal networks such as those supporting face-selective and voice-selective circuitries. However, through pressure to integrate face and voice information for individual recognition (38) and communication (39), phylogenetic and ontogenetic experience may generate privileged links between the two systems, due to shared functional goals. Our findings, together with the evidence of a right dominance for face and voice identification, suggest that such privileged links may be nested in the right hemisphere early during human brain development and be particularly susceptible to functional reorganization following early auditory deprivation. Although overall visual responses were below baseline (deactivation) in the right TVA during visual processing in the hearing groups, a non-significant trend for a larger response to faces versus houses (Fig. 1D) as well as a relatively weak face identity adaptation effect were observed. These results may relate to recent evidence showing both visual unimodal and audio-visual bimodal neuronal subpopulations within early voice sensitive regions in the right hemisphere of hearing macaques (35). It is therefore plausible that in early absence of acoustic information, the brain reorganizes itself by building on existing cross-modal inputs in right temporal regions.

The neuronal mechanisms underlying cross-modal plasticity have yet to be elucidated in humans, although unmasking of existing synapses, ingrowth of existing and rewiring of new connections are thought to support cortical reorganization (24). Our observation that increased feed-forward effective connectivity from early extra-striate visual regions primarily sustains the face-selective response detected in right dTFA provides supporting evidence in favor of the view that cross-modal plasticity could occur early in the hierarchy of brain areas and that reorganization of long-range connections between sensory cortices may play a key role in functionally selective cross-modal plasticity. This is consistent with recent evidence that cross-modal visual recruitment of the pSTS was associated with increased functional connectivity with the calcarine cortex in the Deaf, although the directionality of the effect was undetermined (40). The hypothesis that the auditory cortex participates in early sensory/perceptual processing following early auditory deprivation, in contrast with previous assumptions that such recruitment manifests only for late and higher-level cognitive process (41, 42), also find support in our MEG finding that face-selective response occurs at about 196ms in right dTFA. Since at least 150ms of information accumulation is necessary for high-level individuation of faces in the cortex (22), this suggests that the face-selective response in right dTFA occurs immediately after the initial perceptual encoding of face identity. Similar to our findings, auditory-driven activity in reorganized visual cortex in congenitally blind individuals was also better explained by direct connections with primary auditory cortex (43), whereas it depended more on feedback inputs from high-level parietal regions in late-onset blindness (43). The crucial role of developmental periods of auditory deprivation in shaping the reorganization of long-range cortico-cortical connections remains, however, to be determined.

In summary, these findings confirm that cross-modal inputs might remap selectively onto regions sharing common functional purposes in the auditory domain in early deaf people. Our findings also indicate that reorganization of direct long-range connections between auditory and early visual regions may serve as a prominent neuronal mechanism for functionally selective cross-modal colonization of specific auditory regions in the deaf. These observations are clinically relevant since they might contribute informing the evaluation of potential compensatory forms of cross-modal plasticity and their role in person information processing following early and prolonged sensory deprivation. Moreover, assessing the presence of such functionally specific crossmodal reorganizations may prove important when considering auditory reafferentation via cochlear implant (1).

## Materials and Methods

The research presented in this article was approved by Scientific Committee of CIMeC and the Committee for Research Ethics of the University of Trento. Informed consent was obtained from each participant in agreement with the ethical principles for medical research involving human subjects (Declaration of Helsinki; WMA) and the Italian law on individual privacy (D.l. 196/2003).

### Participants

Fifteen deaf subjects, 16 hearing subjects and 15 hearing LIS users participated in the fMRI study. Seventeen deaf and 14 hearing subjects successively participated in the MEG study; since 3 out of 15 deaf participants who were included in the fMRI study could not return to the laboratory and take part in the MEG study, an additional group of 5 deaf participants was recruited for the MEG experiment only. The three groups participating in the fMRI experiment were matched for age, gender, handedness (44) and non-verbal IQ(45) as were the deaf and hearing groups included in the MEG experiment (Table 2). No participants had reported neurological or psychiatric history and all had normal or corrected-to-normal vision. Information on hearing status, history of hearing loss and use of hearing aids were collected in deaf participants through a structured questionnaire (Table S1). Similarly, information about sign language age of acquisition, duration of exposure and frequency of use was documented in both the deaf and hearing-LIS group and no significant differences were observed between the two groups (Tables 2 and Table S2).

**Table 2.** Demographics, behavioral performances and Italian Sign Language aspects of the 46 subjects participating in the fMRI experiment and of the 31 subjects participating in the MEG experiment.

| Demographics<br>Cognitive Tests | Hearing Controls<br>(n =16) | Hearing-LIS<br>(n=15) | Deaf<br>(n=15) | Statistics |
| --- | --- | --- | --- | --- |
| Mean Age<br>Years (s.d.) | 30.81 (5.19) | 34.06(5.96) | 32.26(7.23) | F-test=1.079<br>P-value = 0.349 |
| Gender (M/F) | 8/8 | 5/10 | 7/8 | $\chi^2 = 0.97$<br>P-value = 0.617 |
| Hand Preference<br>(% Right/left ratio) | 71.36(48.12) | 61.88(50.82) | 58.63(52.09) | K-W test = 1.15<br>P-value = 0.564 |
| IQ<br>Mean estimate (s.d.) | 122.75(8.95) | 124.76(5.61) | 120.23(9.71) | F-test=0.983<br>P-value = 0.384 |
| Composite<br>Face Recognition<br>z-scores (s.d) | -0.009(1.51)** | 1.709(1.40) | 1.704(1.75) | F-test=5.261<br>P-value = 0.010 |
| LIS Exposure<br>Years (s.d.) | -- | 25.03(13.84) | 21.35(9.86) | t-test=-0.079<br>P-value =0.431 |
| LIS Acquisition<br>Years (s.d.) | -- | 11.42(8.95) | 9.033(11.91) | M-W.U-test = 116.5<br>P-value =0.374 |
| LIS Frequency<br>Percent time/year (s.d.) | -- | 70.69(44.00) | 84.80(26.09) | M-W. U-test = 97<br>p-value=0.441 |
| Demographics<br>Cognitive Tests | Hearing Controls<br>(n =14) | -- | Deaf<br>(n =17) | Statistics |
| Mean Age<br>Years (s.d.) | 30.64 (5.62) |  | 35.47(8.59) | t-test=1.805<br>P-value = 0.082 |
| Gender (M/F) | 6/8 | | 7/10 | $\chi^2 = 0.009$<br>P-value < 0.925 |
| Hand Preference<br>(% Right/left ratio) | 74.75(33.45) |  | 78.87(24.40) | M-W.U-test = 110<br>P-value = 1 |
| IQ<br>Mean estimate (s.d.) | 123.4(8.41) |  | 117.8(12.09) | M-W.U-test = 85.5<br>P-value = 0.567 |
| Benton Face Recognition<br>Task<br>z-scores (s.d) | -0.539 (0.97)* |  | 0.430(0.96) | t-test=2.594<br>P-value = 0.016 |

(\*\*) in Deaf versus Hearing Controls and in Hearing-LIS versus Hearing Controls ( $P < 0.025$ ); (\*) in Deaf versus Hearing Controls ( $P = 0.012$ ); Abbreviations: s.d., standard deviation.

### Experimental design: Behavioral Testing

The long version of the Benton Facial Recognition Test (BFRT)(46) and a delayed face recognition test (DFRT), developed specifically for the present study, were used to obtain a composite measure of individual face identity processing in each group(47). The DFRT was administered 10 to 15 minutes after completion of the face localizer fMRI experiment and presented the subjects with 20 images for each category (faces and houses) half of which they had previously seen in the scanner (see dedicated section below). Subjects were instructed to indicate whether they thought they had previously seen the given image.

### Experimental design: fMRI Face Localizer

The Face Localizer task was administered to the three groups (hearing, hearing-LIS, deaf; see Table S2). Two categories of stimuli were used: images of Faces and Houses equated for low-level properties. The Face condition consisted of 20 pictures of static faces with neutral expression and in a frontal view (Radboud Faces database) equally representing male and female individuals (10/10). Similarly, the House condition consisted of 20 full-front photographs of different houses. Low-level image properties (mean luminance, contrast and spatial frequencies) were equated across stimuli categories by editing them with the SHINE (48) toolbox for Matlab (Mathworks inc.). A block-designed one-back identity task was implemented in a single run lasting for about 10 minutes (Fig. S5). Participants were presented with 10 blocks of 21s duration for each of the two categories of stimuli. In each block, 20 stimuli of the same condition were presented (1000 ms, ISI 50ms) on a black background screen; in one to three occasions per block, the exact same stimulus was consecutively repeated that the participant had to detect. Blocks were alternated with a resting baseline condition (cross-fixation) of 7 to 9 sec.

### Experimental design: fMRI voice-localizer and fMRI Face-adaptation

For these experiments we adapted two fMRI-design previously validated (Beline et al, 2000; Gentile and Rossion, 2014). See SI for a detailed description.

### fMRI acquisition parameters

For each fMRI experiment, whole-brain images were acquired at the Center for Mind and Brain Sciences (University of Trento) on a 4 Tesla Brucker BioSpin MedSpec head scanner using a standard head coil and gradient echo planar imaging (EPI) sequences. Acquisition parameters for each experiment are reported in Table S3. Both signing and non-signing deaf individuals could communicate through overt speech or by using a forced choice button-press code previously agreed with the experimenters. In addition, a three-dimensional MP-RAGE T1-weighted image of the whole brain was also acquired in each participant to provide detailed anatomy (176 slices; TE = 4.18ms; TR = 2700 ms; FA = 7°, slice thickness = 1mm).

### Behavioral data analysis

We computed a composite measure of face recognition with unit-weighted z-scores of the BFRT and DFRT to provide a more stable measure of the underlying face processing abilities, as well as control for the number of independent comparisons. A detailed description of the composite calculation is reported in the supplemental material.

### Functional MRI data analysis

We analyzed each fMRI dataset using SPM12 (http://www.fil.ion.ucl.ac.uk/spm12) and Matlab R2012b (The Matworks, Inc).

#### Preprocessing of fMRI data

For each subject and for each dataset, the first 4 images were discarded to allow magnetic saturation effects. The remaining images in each dataset (Face Localizer = 270; Voice Localizer = 331; Face-adaptation = 329 x 3 runs) were visually inspected and a first manual co-registration between the individual first EPI volume of each dataset, the corresponding MP-RAGE volume and the T1 Montreal Neurological Institute (MNI) template was performed. Subsequently, in each dataset, the images were corrected for timing differences in slice acquisition, motion corrected (6 parameter affine transformation) and realigned to the mean image of the corresponding sequence. The individual T1 image was segmented in grey and white matter parcellations and the forward deformation field computed. Functional EPI images (3mm isotropic voxels) and the T1 image (1mm isotropic voxels) were normalized to the MNI space using the forward deformation field parameters and data resampled at 2mm isotropic with a 4^th^ degree B-spline interpolation. Finally, the EPI images in each dataset were spatially smoothed with a Gaussian kernel of 6mm full width at half maximum (FWHM).

For each fMRI experiment, first-level (single-subject) analysis used a design matrix including separate regressors for the conditions of interest (see below), plus realignment parameters to account for residual motion artifacts as well as outlier regressors; these regressors referred both to scans with large mean displacement and/or weaker or stronger globals. The regressors of interest were defined by convolving boxcars functions representing the onset and offset of stimulation blocks in each experiment by the canonical hemodynamic response function (HRF). Each design matrix also included a filter at 128s and auto-correlation, which was modeled using an auto-regressive matrix of order 1.

#### fMRI Face Localizer modeling

Two predictors corresponding to face and house images were modeled and the contrast [face > house] was computed for each participant; these contrast images were then further spatially smoothed by a 6mm FWHM prior to group-level analyses. The individual contrast images of the participants were entered in a one-sample *t*-test to localize regions showing face-selective response in each group. Statistical inference was made at a corrected cluster level of *p* < 0.05 FWE (with a standard voxel-level threshold of *p* < 0.001 uncorrected) and a minimum cluster-size of 50. Subsequently, a one-way ANOVA was modeled with the three groups as independent factors and a conjunction analysis (deaf _[Face > House]_ > hearing_[Face > House]_ *conjunction with* deaf _[Face > House]_ > hearing-LIS_[Face > House]_) implemented to test for differences between the deaf and the two hearing groups. For this test, statistical inferences were performed also at 0.05 FWE voxel-corrected over a small spherical volume (25 mm radius) located at the peak coordinates of group-specific response to vocal sound in the left and right STG/STS, respectively, in hearing subjects (Table 1). Consequently, measures of individual response to faces and houses were extracted from the right and left TVA in each participant. To account for inter-individual variability a search-sphere of 10 mm radius was centered at the peak coordinates (x = 63, y = −22, z = −4; x = −60, y = −16, z = 1; MNI) corresponding to the group-maxima for [Vocal > Non-Vocal Sounds] in the hearing group. Additionally, the peak-coordinates search was constrained by the TVA masks generated in our hearing group to exclude extraction from posterior STS/STG associative sub-regions that are known to be also involved in face processing in hearing individuals. Finally, the corresponding beta values were extracted from a 5 mm sphere centered on the selected individual peak coordinates (see also supplemental information). These values were then entered in a repeated measure ANOVA with the two visual conditions as within-subject factor and the three groups as between-group factor.

#### fMRI Face-adaptation modeling

We implemented a GLM with 10 regressors corresponding to the [5 frequencies × same/different] face images and computed the contrast images for the [Same/Different Face versus baseline (cross-fixation)] test at each frequency rate of visual stimulation. In addition, the contrast image [Different versus Same Faces] across frequency rates of stimulation was also computed in each participant; at the group level, these contrast images were entered as independent variables in three one-sample *t*-tests, separately and specifically for each experimental group, in order to evaluate whether discrimination of individual faces elicited the expected responses within the face-selective brain network (voxel significance at p < 0.05 FWE-corrected). Subsequent analyses were restricted to the functionally defined face- and voice-sensitive areas (Voice and Face localizers; see above) from which the individual beta values corresponding to each condition were extracted. The Bonferroni correction was applied to correct for multiple comparisons as appropriate.

#### fROI definition for face-adaptation

In each participant and for each region, this was achieved by: (i) centering a sphere-volume of 10 mm radius at the peak-coordinates reported for the corresponding group, (ii) anatomically constraining the search within the relevant cortical gyrus (e.g. for the right FFA the right fusiform as defined by the Automated Anatomical Labeling atlas in SPM12), and (iii) extracting condition-specific mean beta values from a sphere volume of 5 mm radius (Table S4). The extracted betas were then entered as dependent variables in a series of repeated measures ANOVAs and *t*-tests as reported in the main result session.

### Experimental design: MEG Face Localizer

A Face Localizer task in the MEG was recorded from 14 hearing (age 30.64) and 17 deaf subjects (age 35.47); all participants except for 5 deaf subjects also participated in the fMRI part of the study. Participants viewed the stimulus at a distance from the screen of 100cm. The images of 40 faces and 40 houses were identical to the ones used in fMRI. Afterfixation period (1000-1500ms) the visual image was presented for 600ms. Participants were instructed to press a button whenever an image was presented twice in a row (oddball). Catch trials (~11%) were excluded from subsequent analysis. The images were presented in a pseudo-randomized fashion and in three consecutive blocks. Every stimulus was repeated three times, adding up to a total number of 120 trials per condition.

### MEG data acquisition

MEG was recorded continuously on a 102 triple sensor (two gradiometer, and one magnetometer) whole-head system (Elekta Neuromag, Helsinki Finland). Data was acquired with a sampling rate of 1kHz and an online band pass filter between 0.1 and 330 Hz. Individual headshapes were recorded using a Polhemus FASTRAK 3D digitizer. The head position was measured continuously using five localization coils (forehead, mastoids). For improved source reconstruction individual structural MR images were acquired on a 4T scanner (Bruker Biospin Ettlingen, Germany).

### MEG data analysis

#### Preprocessing

The data preprocessing and analysis was performed using the open-source toolbox fieldtrip (49) as well as custom Matlab codes. The continuous data was filtered (high-pass Butterworth filter at 1Hz; DFT filter at 50,100, and 150Hz) and downsampled to 500 Hz to facilitate computational efficiency. Analyses were performed on the gradiometer data. The filtered continuous data was epoched around the events of interest, and inspected visually for muscle and jump artifacts. Remaining ocular and cardiac artifacts were removed from the data using extended infomax independent component analysis (ICA) with a weight change stop criterion of 10^−7^. Finally, a pre-stimulus baseline of 150ms was applied to the cleaned epochs.

#### Sensor-space analysis

Sensor-space analysis was performed across groups prior to source-space analyses. The cleaned data were low-pass filtered at 30 Hz and averaged separately across face and house trials. Statistical comparisons between the two conditions were performed using a cluster permutation approach in space (sensors) and time (50) in a time window between 100 and 300ms after stimulus onset. Adjacent points in time and space exceeding a predefined threshold (*p*<.05) were grouped into one or multiple clusters, and the summed cluster t-values were compared against a permutation distribution. The permutation distribution was generated by randomly reassigning condition membership for each participant (1000 iterations), and computing the maximum cluster mass on each iteration. This approach reliably controls for multiple comparisons at the cluster-level. The time period with the strongest difference between faces and houses was used to guide subsequent source analysis. To illustrate global energy fluctuations during the perception of faces and houses global field power (GFP) was computed as the root mean square (RMS) of the averaged response to the two stimulus types across sensors.

#### Source-space analysis

Functional data was co-registered with the individual subject MRI using anatomical landmarks (pre-auricular points and nasion), and the digitized headshape to create a realistic single-shell headmodel. When no individual structural MRI was available (5 participants), a model of the individual anatomy was created by warping an MNI template brain to the individual subject’s headshape. Broadband source power was projected onto a 3-dimensional grid (8mm spacing) using linear constrained minimum variance (LCMV) beamforming. To ensure stable und unbiased filter coefficients, a common filter was computed from the average covariance matrix across conditions between 0 and 500ms after stimulus onset. Whole-brain statistics were performed using a two-step procedure. First, independent-samples t-tests were computed for the difference between face and house trials by permuting condition membership (1000 iterations). The resulting statistical T-maps were converted to Z-maps for subsequent group analysis. Finally, second-level group statistics were performed using statistical non-parametric mapping (SnPM) and family wise error (FWE) correction at *p*<.05 was applied to correct for multiple comparisons. To further explore the time course of face processing in FFA and dTFA for the early deaf participants virtual sensors were computed on the 40Hz low-pass filtered data using an LCMV beamformer at the FFA and TVA/dTFA locations of interest identified in the whole-brain analysis. As the polarity of the signal in source space is arbitrary, we computed the absolute for all virtual sensor time-series. A baseline correction of 150ms pre-stimulus was applied to the data.

### fMRI functional connectivity analysis

Task-dependent contributions of the right dTFA and TVA to brain face-selective responses elsewhere were assessed in the deaf and in the hearing groups, respectively, by implementing a Psycho-Physiological Interactions (PPI) analysis (51) on the fMRI face localizer dataset. The individual time series for the right TVA/dTFA were obtained by extracting the first principal component from all raw voxel time series in a sphere (radius = 5 mm) centered on the peak-coordinates of the subject-specific activation in this region (i.e. face-selective responses in deaf subjects and voice-selective responses in hearing subjects and hearing LIS users). After individual time series had been mean-corrected and high-pass filtered to remove low-frequency signal drifts, a PPI term was computed as the element-by-element product of the TVA/dTFA time series and a vector coding for the main effect of task (1 for face presentation, −1 for house presentation). Subsequently, a GLM was implemented including the PPI term, region-specific time-series, main effect of task vector, moving parameters and outlier scans vector as model regressors. The contrast image corresponding to the positive-tailed one-sample *t*-test over the PPI regressor was computed in order to isolate brain regions receiving stronger contextual influences from the right TVA/dTFA during face processing compared to house processing. The opposite, i.e. stronger influences during house processing, was achieved by computing the contrast image for the negative-tailed one-sample *t*-test over the same PPI regressor. These subject-specific contrast images were spatially smoothed by a 6mm FWHM prior submission to subsequent statistical analyses. For each group, individually smoothed contrast images were entered as the dependent variable in a one-sample *t*-test to isolate regions showing face-specific increased functional connectivity with the right TVA/dTFA. Finally, individual contrast images were also entered as the dependent variable in two one-way ANOVAs, one for face and one for house responses, with the three groups as between-subject factor to detect differences in functional connectivity from TVA/dTFA between groups. For each test, statistical inferences were made at corrected cluster level of *p* < 0.05 FWE (with a standard voxel-level threshold of *p* < 0.001 uncorrected) with a minimum size of 50 voxels.

### Effective Connectivity Analysis

Dynamic Causal Modeling (DCM)(52), a hypothesis-driven analytical approach, was used to characterize the causality between the activity recorded in the set of regions that showed increased functional connectivity with the right dTFA in the deaf group during face compared to house processing. To this purpose, our model space was operationalized based on three neurobiologically plausible and sufficient alternatives: (i) face-selective response in right dTFA is supported by increased connectivity modulation directly from right V2/V3, (ii) face selective response in the right dTFA is supported indirectly by increased connectivity modulation from right FFA or, (iii) face selective response in the right dTFA is supported indirectly by increased connectivity modulation from right pSTS. DCM models can only be used for investigating brain responses that present a relation to the experimental design and can be observed in each individual included in the investigation(52). Since no temporal activation was detected for face and house processing in hearing subjects and hearing LIS users, these groups were not included in the DCM analysis. For a detailed description of DCMs see supplemental information.

The three DCMs were fitted with the data from each of the 15 deaf participants; this resulted in 45 fitted DCMs and corresponding log-evidence and posterior parameters estimates. Subsequently, random-effect Bayesian Model Selection (53) was applied to the estimated evidence for each model to compute the ‘exceedance probability’. This is the probability of each specific model to better explain the observed activations compared to any other model.

## Acknowledgments

This work was supported by the ‘Società Mente e Cervello’ of the Center for Mind/Brain Sciences (University of Trento; S.B., F.B., O.C.). The authors wish to thank all the deaf people and the hearing sign language users, who participated in this research, for their collaboration and support throughout the completion of the study.

## References

1. Heimler B, Weisz N, Collignon O (2014) Revisiting the adaptive and maladaptive effects of crossmodal plasticity. Neuroscience 283:44–63.

2. Finney EM, Fine I, Dobkins KR (2001) Visual stimuli activate auditory cortex in the deaf. Nat Neurosci 4(12):1171–1173.

3. Karns CM, Dow MW, Neville HJ (2012) Altered cross-modal processing in the primary auditory cortex of congenitally deaf adults: a visual-somatosensory fMRI study with a double-flash illusion. J Neurosci 32(28):9626–9638.

4. Lomber SG, Meredith MA, Kral A (2010) Cross-modal plasticity in specific auditory cortices underlies visual compensations in the deaf. Nat Neurosci 13(11):1421–1427.

5. Bola Ł, et al. (2017) Task-specific reorganization of the auditory cortex in deaf humans. Proc Natl Acad Sci Early Edit: 1–10.

6. Pavani F, Bottari D (2012) Visual Abilities in Individuals with Profound Deafness A Critical Review. The Neural Bases of Multisensory Processes, eds Murray MM, Wallace MT (CRC Press, Boca Raton (FL)). Available at: http://file//localhost(null).

7. Lee H-J, et al. (2007) Cortical activity at rest predicts cochlear implantation outcome. Cereb cortex (New York, NY 1991) 17(4):909–917.

8. Bettger J, Emmorey K, McCullough S, Bellugi U (2004) Enhanced facial discrimination: effects of experience with American sign language. J Deaf Stud Deaf Educ 2(4):223–233.

9. Rouger J, Lagleyre S, Démonet JF, Barone P (2012) Evolution of crossmodal reorganization of the voice area in cochlear-implanted deaf patients. Hum brain… 33(8):1929–1940.

10. Stropahl M, et al. (2015) Cross-modal reorganization in cochlear implant users: Auditory cortex contributes to visual face processing. Neuroimage 121:159–170.

11. Belin P, Zatorre RJ, Lafaille P, Ahad P, Pike B (2000) Voice-selective areas in human auditory cortex. Nature 403(6767):309–312.

12. Ghazanfar AA, Maier JX, Hoffman KL, Logothetis NK (2005) Multisensory integration of dynamic faces and voices in rhesus monkey auditory cortex. J Neurosci 25(20):5004–12.

13. Kanwisher N, McDermott J, Chun MM (1997) The fusiform face area: a module in human extrastriate cortex specialized for face perception. J Neurosci 17(11):4302–11.

14. von Kriegstein K, Kleinschmidt A, Sterzer P, Giraud A-L (2005) Interaction of face and voice areas during speaker recognition. J Cogn Neurosci 17(3):367–76.

15. Blank H, Anwander A, von Kriegstein K (2011) Direct structural connections between voice- and face-recognition areas. J Neurosci 31(36):12906–12915.

16. Grill-Spector K, Malach R (2001) fMR-adaptation: a tool for studying the functional properties of human cortical neurons. Acta Psychol (Amst) 107(1–3):293–321.

17. Jacques C, et al. (2016) Corresponding ECoG and fMRI category-selective signals in human ventral temporal cortex. Neuropsychologia 83:14–28.

18. Benton AL, Van Allen MW (1968) Impairment in Facial Recognition in Patients with Cerebral Disease. Cortex 4(4):344–358.

19. Hildebrandt A, Sommer W, Herzmann G, Wilhelm O (2010) Structural invariance and age-related performance differences in face cognition. Psychol Aging 25(4):794–810.

20. Arnold P, Murray C (1998) Memory for faces and objects by deaf and hearing signers and hearing nonsigners. J Psycholinguist Res 27(4):481–497.

21. Gentile F, Rossion B (2014) Temporal frequency tuning of cortical face-sensitive areas for individual face perception. Neuroimage 90:256–265.

22. Alonso-Prieto E, Belle G Van, Liu-Shuang J, Norcia AM, Rossion B (2013) The 6Hz fundamental stimulation frequency rate for individual face discrimination in the right occipito-temporal cortex. Neuropsychologia 51(13):2863–2875.

23. Halgren E, Raij T, Marinkovic K, Jousmäki V, Hari R (2000) Cognitive response profile of the human fusiform face area as determined by MEG. Cereb Cortex 10(1):69–81.

24. Bavelier D, Neville HJ (2002) Cross-modal plasticity: where and how? Nat Rev Neurosci 3(6):443–452.

25. Lohse M, et al. (2016) Effective Connectivity from Early Visual Cortex to Posterior Occipitotemporal Face Areas Supports Face Selectivity and Predicts Developmental Prosopagnosia. J Neurosci 36(13):3821–3828.

26. Pernet CR, et al. (2015) The human voice areas: Spatial organization and inter-individual variability in temporal and extra-temporal cortices. Neuroimage 119:164–174.

27. Dormal G, Collignon O (2011) Functional selectivity in sensory-deprived cortices. J Neurophysiol. Available at: http://jn.physiology.org/content/105/6/2627.abstract.

28. Latinus M, McAleer P, Bestelmeyer PEG, Belin P (2013) Norm-based coding of voice identity in human auditory cortex. Curr Biol 23(12):1075–1080.

29. Reich L, Szwed M, Cohen L, Amedi A (2011) A Ventral Visual Stream Reading Center Independent of Visual Experience. Curr Biol 21(5):363–368.

30. Collignon O, et al. (2011) Functional specialization for auditory-spatial processing in the occipital cortex of congenitally blind humans. Proc Natl Acad Sci U S A 108(11):4435–4440.

31. Collignon O, Voss P, Lassonde M, Lepore F (2009) Cross-modal plasticity for the spatial processing of sounds in visually deprived subjects. Exp brain Res Exp Hirnforsch Expérimentation cérébrale 192(3):343–358.

32. Pascual-Leone A, Hamilton R (2001) The metamodal organization of the brain. Progress in Brain Research, pp 427–445.

33. Amedi a, Malach R, Hendler T, Peled S, Zohary E (2001) Visuo-haptic object-related activation in the ventral visual pathway. Nat Neurosci 4(3):324–330.

34. Yovel G, Belin P (2013) A unified coding strategy for processing faces and voices. Trends Cogn Sci 17(6):263–271.

35. Perrodin C, Kayser C, Logothetis NK, Petkov CI (2014) Auditory and visual modulation of temporal lobe neurons in voice-sensitive and association cortices. J Neurosci 34(7):2524–2537.

36. Belin P, Zatorre RJ (2003) Adaptation to speaker’s voice in right anterior temporal lobe. Neuroreport 14(16):2105–2109.

37. MacSweeney M, Capek CM, Campbell R, Woll B (2008) The signing brain: the neurobiology of sign language. Trends Cogn Sci 12(11):432–440.

38. Sheehan MJ, Nachman MW (2014) Morphological and population genomic evidence that human faces have evolved to signal individual identity. Nat Commun 5:4800.

39. Ghazanfar A a, Logothetis NK (2003) Neuroperception: facial expressions linked to monkey calls. Nature 423(6943):937–938.

40. Shiell MM, Champoux F, Zatorre RJ (2014) Reorganization of Auditory Cortex in Early-deaf People: Functional Connectivity and Relationship to Hearing Aid Use. J Cogn Neurosci 21:1–14.

41. Leonard MK, et al. (2012) Signed words in the congenitally deaf evoke typical late lexicosemantic responses with no early visual responses in left superior temporal cortex. J Neurosci 32(28):9700–9705.

42. Ding H, et al. (2015) Cross-modal activation of auditory regions during visuo-spatial working memory in early deafness. Brain 138(9):2750–2765.

43. Collignon O, et al. (2013) Impact of blindness onset on the functional organization and the connectivity of the occipital cortex. Brain 136(9):2769–2783.

44. Oldfield RC (1971) The assessment and analysis of handedness: The Edinburgh inventory. Neuropsychologia 9(1):97–113.

45. Raven J, Raven JC, Court J (1998) Manual for Raven’s progressive matrices and vocabulary scales Available at: http://books.google.es/books?id=YrvAAQAACAAJ.

46. Benton A, Hamsher K (1983) Multilingual Aphasia Examinationle doi:10.1076/clin.15.1.13.1911.

47. Ackerman PL, Cianciolo a T (2000) Cognitive, perceptual-speed, and psychomotor determinants of individual differences during skill acquisition. J Exp Psychol Appl 6(4):259–290.

48. Willenbockel V, et al. (2010) Controlling low-level image properties: the SHINE toolbox. Behav Res Methods 42(3):671–684.

49. Oostenveld R, Fries P, Maris E, Schoffelen JM (2011) FieldTrip: Open source software for advanced analysis of MEG, EEG, and invasive electrophysiological data. Comput Intell Neurosci 2011. doi:10.1155/2011/156869.

50. Maris E, Oostenveld R (2007) Nonparametric statistical testing of EEG- and MEG-data. J Neurosci Methods 164(1):177–190.

51. Friston KJ, et al. (1997) Psychophysiological and modulatory interactions in neuroimaging. Neuroimage 6(3):218–229.

52. Stephan KE (2006) Dynamic causal models of neural system dynamics: current state and future extensions. 1–16.

53. Stephan KE, Penny WD, Daunizeau J, Moran RJ, Friston KJ (2009) Bayesian model selection for group studies. Neuroimage 46(4):1004–1017.

